## Supplementary material for "Deep learning generates custom-made logistic regression models for explaining how breast cancer subtypes are classified": S1 Fig. Paper overview.

Quick look


### Introduction

### Materials and Methods

- ✓ Point-wise linear models
- ✓ Datasets
- Feature importance calculation method
- Deep enrichment analysis method
- Evaluation methodology

### Results

- Prediction performance
- ✓ Feature importance analysis
- ✓ Enrichment analysis

### Discussion

### Conclusion
